# Transition into inflammatory cancer-associated adipocytes in breast cancer microenvironment requires microRNA regulatory mechanism

**DOI:** 10.1101/089243

**Authors:** Jiwoo Lee, Han Suk Ryu, Bok Sil Hong, Han-Byoel Lee, Minju Lee, In Ae Park, Jisun Kim, Wonshik Han, Dong-Young Noh, Hyeong-Gon Moon

## Abstract

**Introduction:** The role of adipocytes in cancer microenvironment has gained focus during the recent years. However, the characteristics of the cancer-associated adipocytes (CAA) in human breast cancer tissues and the underlying regulatory mechanism are not clearly understood.

**Method:** We reviewed pathology specimens of breast cancer patients to understand the morphologic characteristics of CAA, and profiled the mRNA and miRNA expression of CAA by using indirect co-culture system *in vitro*.

**Results:** The CAAs in human breast cancers showed heterogeneous topographic relationship with breast cancer cells within the breast microenvironment. The CAAs exhibited the characteristics of de-differentiation determined by their microscopic appearance and the expression levels of adipogenic markers. Additionally, the 3T3-L1 adipocytes co-cultured with breast cancer cells showed up-regulation of inflammation-related genes including *Il6* and *Ptx3*. The up-regulation of IL6 in CAA was further observed in human breast cancer tissues. miRNA array of co-cultured 3T3-L1 cells showed increased expression of mmu-miR-5112 which may target *Cpeb1*. *Cpeb1* is a negative regulator of *Il6*. The suppressive role of mmu-miR-5112 was confirmed by dual luciferase reporter assay, and mmu-miR-5112-treated adipocytes showed up-regulation of *Il6*. The transition of adipocytes into more inflammatory CAA resulted in proliferation-promoting effect in ER positive breast cancer cells such as MCF7 and ZR-75-1 but not in ER negative cells.

**Conclusion:** In this study, we have determined the de-differentiated and inflammatory natures of CAA in breast cancer microenvironment. Additionally, we propose a miRNA-based regulatory mechanism underlying the process of acquiring inflammatory phenotypes in CAA.

## INTRODUCTION

The association between the degree of obesity and the risk of breast cancer incidence has long been recognized. The proposed mechanisms relating the adipose tissue and the breast cancer risk are mostly explained by endocrine and systemic effects of the adipokines and sex hormones.[1] However, recent studies have focused on the potential roles of adipocytes in the breast cancer microenvironment that may modulate the growth of the cancer cells.[2,3] We have previously reported the poor treatment outcome of underweight breast cancer patients by analyzing a large dataset of Korean breast cancer patients.[4] Interestingly, the worse outcome of underweight patients was most pronounced for the locoregional recurrence rather than distant metastasis suggesting a potential local interaction between adipose tissue and breast cancer progression.

Studies have shown that the adipocytes in the vicinity of the cancer cells, often named as ‘cancer-associated adipocytes (CAA)’, show diverse molecular characteristics that can enhance the ability of cancer cell invasion and aid the cancer cell proliferation via conferring metabolic advantages.[5-7] However, studies have conflicting results on the differentiation lineage of the CAA,[7-9,6,10] and the regulatory mechanisms of the CAA transition is not clearly understood. In this study, we aimed to characterize the CAA in human breast cancer tissues and to explore the regulatory mechanism of the CAA transition.

## MATERIALS AND METHODS

### Pathologic examination of the adipocytes within the breast cancer specimens

Hematoxylin-eosin stained slides of 55 consecutive breast cancer specimens were reviewed to observe the characteristics of the adipocytes. For each patient, representative images showing adipocytes in the vicinity of the epithelial cancer cells and adipocytes around the normal mammary glandular tissue were obtained and the cell sizes were measured with ImageJ.[11] The collection of the clinical and pathologic data from the breast cancer patients was approved by the institutional IRB (1208-046-421) and all procedures were done in accordance with the Declaration of Helsinki.

### Adipogenic differentiation of 3T3-L1 cells and indirect co-culture with cancer cells

3T3-L1 cell line was purchased from ATCC (Manassas, Virginia) and were cultured in DMEM supplemented with 10% bovine serum at 37°C in 5% CO2. 3T3-L1 cells were grown to confluency and induced with adipogenesis media (DMEM with 10% FBS, 10μg/ml insulin, 0.5mM IBMX and 1μM dexamethasone). After 2 days, the media was changed with DMEM with 10% FBS and 10μg/ml insulin and maintained for 4 days. After 8 days of differentiation with DMEM with 10% FBS, the cells were used for further experiments. Breast cancer cell lines used for the co-culture study were MCF7, ZR75-1, T47D, MDA-MB231, and Hs578T. MCF7, MDA-MB231, and Hs578T cells were cultured in DMEM supplemented with 10% FBS, 100 units/ml penicillin, and 100μg/ml streptomycin. ZR75-1 and T47D cells were cultured in RPMI media with same supplements. For the indirect co-culture of the differentiated adipocytes and breast cancer cell lines, we used transwell plate with 0.4μm micron inserts (BD Biosciences, San Jose, CA). Differentiated adipocytes were cultured in the bottom chamber of the transwell co-culture system with or without the breast cancer cells seeded in the top inserts. After the co-culture for certain periods, the adipocytes at the bottom wells were collected and used for further analysis.

### mRNA and miRNA expression microarray

For the quality control, RNA purity and integrity were evaluated by OD 260/280 ratio, and analyzed by Agilent 2100 Bioanalyzer (Agilent Technologies, Palo Alto, USA). For gene expression analysis, the Affymetrix Whole transcript Expression array process was executed according to the manufacturer’s protocol (GeneChip Whole Transcript PLUS reagent Kit). cDNA was synthesized using the GeneChip WT (Whole Transcript) Amplification kit as described by the manufacturer. The sense cDNA was then fragmented and biotin-labeled with TdT (terminal deoxynucleotidyl transferase) using the GeneChip WT Terminal labeling kit. Approximately 5.5 μg of labeled DNA target was hybridized to the Affymetrix GeneChip Human 2.0 ST Array at 45°C for 16hour. Hybridized arrays were washed and stained on a GeneChip Fluidics Station 450 and scanned on a GCS3000 Scanner (Affymetrix). Raw data were extracted automatically in Affymetrix data extraction protocol using the software provided by Affymetrix GeneChip^®^ Command Console^®^ Software (AGCC). After importing CEL files, the data were summarized and normalized with robust multi-average (RMA) method implemented in Affymetrix^®^ Expression Console(tm) Software (EC). We exported the result with gene level RMA analysis and perfomed the differentially expressed gene (DEG) analysis.

For miRNA analysis, the Affymetrix Genechip miRNA array process was executed according to the manufacturer’s protocol. 1μg RNA samples were labeled with the FlashTag(tm) Biotin RNA Labeling Kit (Genisphere, Hatfield, PA, USA). The labeled RNA was quantified, fractionated and hybridized to the miRNA microarray according to the standard procedures provided by the manufacture. The labeled RNA was heated to 99°C for 5 minutes and then to 45°C for 5 minutes. RNA-array hybridization was performed with agitation at 60 rotations per minute for 16–18 hours at 48°C on an Affymetrix^®^ 450 Fluidics Station. The chips were washed and stained using a Genechip Fluidics Station 450 (Affymetrix, Santa Clara, CA). The chips were then scanned with an Affymetrix GeneChip Scanner 3000 (Affymetrix, Santa Clara, CA). Signal values were computed using the Affymetrix^®^ GeneChip(tm) Command Console software. Raw data were extracted automatically in Affymetrix data extraction protocol using the software provided by Affymetrix GeneChip^®^ Command Console^®^ Software. The CEL files import, miRNA level RMA+DABG-All analysis and result export using Affymetrix^®^ Expression Console(tm) Software. Array data were filtered by probes annotated species.

Differential expression was analyzed using independent t-test and Benjamini-Hochberg algorithm. Gene-Enrichment and Functional Annotation analysis for significant probe list was performed using DAVID (http://david.abcc.ncifcrf.gov/). All Statistical test and visualization of differentially expressed genes was conducted using R statistical language v. 3.0.2. (www.r-project.org).

### Enzyme-linked immunosorbent assay (ELISA)

After 3 days of co-culture with cancer cells, the inserts containing cancer cells were removed and the remaining adipocytes were washed with PBS and were cultured for additional 2 days. The media was collected and the level of IL-6 was determined by enzyme-linked immunosorbent assay. Mouse IL-6 antibody was purchased from eBioscience (CA, USA). After 16 hr of incubation at 4°C, 100μl amount of media was aspirated from each well. Biotin-conjugated IL-6 was added to the wells and incubated for 1hr at room temperature. After washing off the unbound antibody-enzyme conjugates, the substrate solution was then added to each well and incubated for 30 min before measuring the absorbance at the wavelength of 450nm.

### Western blot, RT-PCR, and Taqman PCR

After 3 days of co-culture with cancer cells, adipocytes were lysed with RIPA buffer (Thermo scientific, Korea). After incubation on ice for 30 minutes, cellular debris was removed by centrifugation for 10 minutes at 4°C. 10 μg of proteins were separated by SDS-PAGE and then transferred to a polyvinylidene difluoride membrane. After blocking with 5% skim milk, the membranes were probed with an FABP4 antibody (abcam). Blots were developed with an enhanced chemiluminescence Western blotting detection system (Amersham, Buckinghamshire, UK).

For RT-PCR experiments, total RNA from cultured cells was isolated using TRIzol reagent (Life Technologies, Gaithersburg, MD) and was quantified by Nanodrop. 1 ug of total RNAs were reverse-transcribed to cDNA using PrimeScript reverse transcriptase (Takara, Japan) The information of the primers used for this study are shown in the Supplementary file 1. All reaction products were analyzed after 28–35 amplification cycles and expression levels were normalized to the values of Gapdh.

For TaqMan PCR, total RNA was extracted from cells using TRIzol reagent (Favorgen Biotech, Taiwan) according to the manufacturer’s instructions. RNA was converted to cDNA using a TaqMan MicroRNA Reverse Transcription Kit (Applied Biosystems, Foster City, CA). Real-time PCR was performed with TaqMan Universal PCR Master Mix on an Applied Biosystems 7500 System (Applied Biosystems, Foster City, CA). The miRs were normalized to the level of snoRNA412 and the relative expression levels of genes were calculated using the 2-ΔΔ Ct method. All reactions were performed in triplicates.

### Luciferase assay

CPEB1 3’-UTR containing the predicted binding site sequences were obtained by annealing sense and antisense strand or amplifying the genomic DNA from 3T3-L1 cells. The sequences were inserted into XhoI and NotI sites in psiCHECK2 dual luciferase reporter plasmid purchased from Promega ((Madison, WI).

HEK 293FT cells were plated in 24-well plates in DMEM supplemented with 10% fetal bovine serum at 37 °C with 5% CO2. When the cells reached 70–80% confluence, 20 ng of psiCHECK-2 recombination vector and 20 nM of miR-5112 mimics or Negative Control were cotransfected with RNAiMAX (Invitrogen, Carlsbad, CA). After 40 h, the cells were washed with PBS and reseeded in 96-well plates at a density of 5 × 103 cells per well with three replicates. Firefly and Renilla luciferase activities were measured with the Dual-Glo luciferase system (Promega, Madison, WI).

### Proliferation assay

Proliferation was measured using CellTiter-Glo Luminescent Cell viability Assay (Promega, Madison, WI), according to the manufacturer’s instructions. Briefly, after co-culture with adipocytes, cancer cells in the transwell were washed with PBS, trypsinized and reseeded in 96 well plates. Measure the luminescence using microplate luminometer.

## RESULTS

### De-differentiated characteristics of CAAs in breast cancer

First, we assessed the microscopic appearance of CAAs in human breast cancer specimens by investigating samples from 55 consecutive breast cancer patients who underwent curative resection. The CAA, located at the invading front of cancer cells, showed smaller cell size (Fig 1a and 1d), and carried substantial heterogeneity in the patterns of surrounding extracellular matrix and their topographic relationship with cancer cells (Figure 1b and 1c). The smaller cell size is a representative morphologic features of the less differentiated adipocytes (Supplementary Figure 1). These findings led us to hypothesize that the observed microscopic features of the CAA could be resulting from a de-differentiation process of mature adipocytes under the influence of the adjacent cancer cells. The possibility of de-differentiation process in CAA has also been suggested by previous studies.[12,13,6] To demonstrate the de-differentiation process of the adipocytes under the influence of breast cancer cells, we used indirect co-culture system for 3T3-L1 preadipocytes and various breast cancer cells. Compared to the adipocyte cultured alone, the adipocytes co-cultured with breast cancer cells showed smaller cell size and reduced lipid drops within the cells (Figure 2a). Also, the markers of differentiated adipocytes, such as C/EBP-α, PPAR-γ, and FABP4, were down-regulated in co-cultured adipocytes (Figure 2b and 2c). These data suggest that cancer cells influence the adjacent adipocytes by paracrine effect and induce the de-differentiation process.

**Figure 1.**
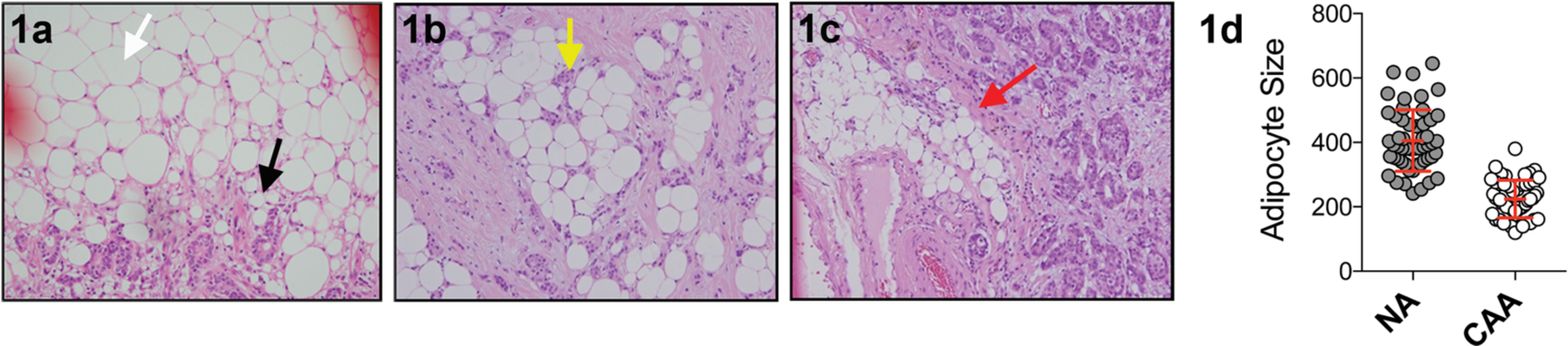
Microscopic appearances of the cancer-associated adipocytes (CAA) in human breast cancer specimens. The CAA (black arrow) showed smaller lipid droplets and hence smaller cell size when compared to normal adipocytes (white arrow) (1a). In some cases, the breast cancer cells were clustered in contact with the CAA (1b, yellow arrow) while other cases showed stromal tissues separating the nests of cancer cells and CAA (1c, red arrow). The cell diameters of CAA and NA (normal adipocyte) are shown in 1d (arbitrary unit, measured by ImageJ).

**Figure 2.**
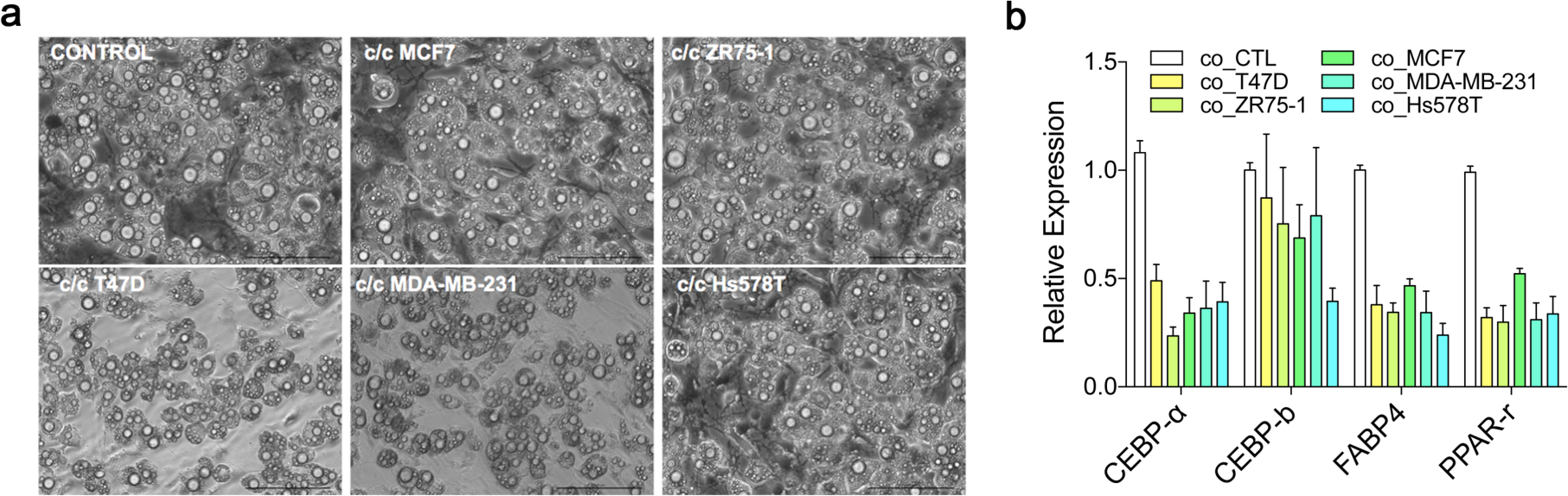
The microscopic appearance and expression characteristics of adipocytes when co-cultured with various breast cancer cells. Adipocytes co-cultured with various cancer cells showed reduction in lipid drops and cell size (a). Genes and proteins involved in the adipocyte differentiation showed patterns of down-regulation in adipocytes co-cultured with breast cancer cells (b and c)

### Effect of CAA on the proliferative capacity of breast cancer cells

To determine the effect of cancer-associated adipocytes on the proliferation of breast cancer cells, we co-cultured the cancer cells with mature adipocytes for three days. After removal of the cancer cells, a new set of cancer cells were introduced to the co-cultured adipocytes and cultured for three days (Figure 3a). As shown in the Figure 3b, the effect of CAA on the proliferation of breast cancer cells varied between the cell lines. The pro-proliferative effect of CAA was most evident in the ER positive cells such as MCF7 and ZR75-1 while no effect was shown for ER negative cells including MDA-MB-231 and Hs578T. Our results suggest that the proliferation promoting effect of CAA, which has been suggested by previous studies [6-8], might be exerted in a subtype-specific manners in breast cancer.

**Figure 3.**
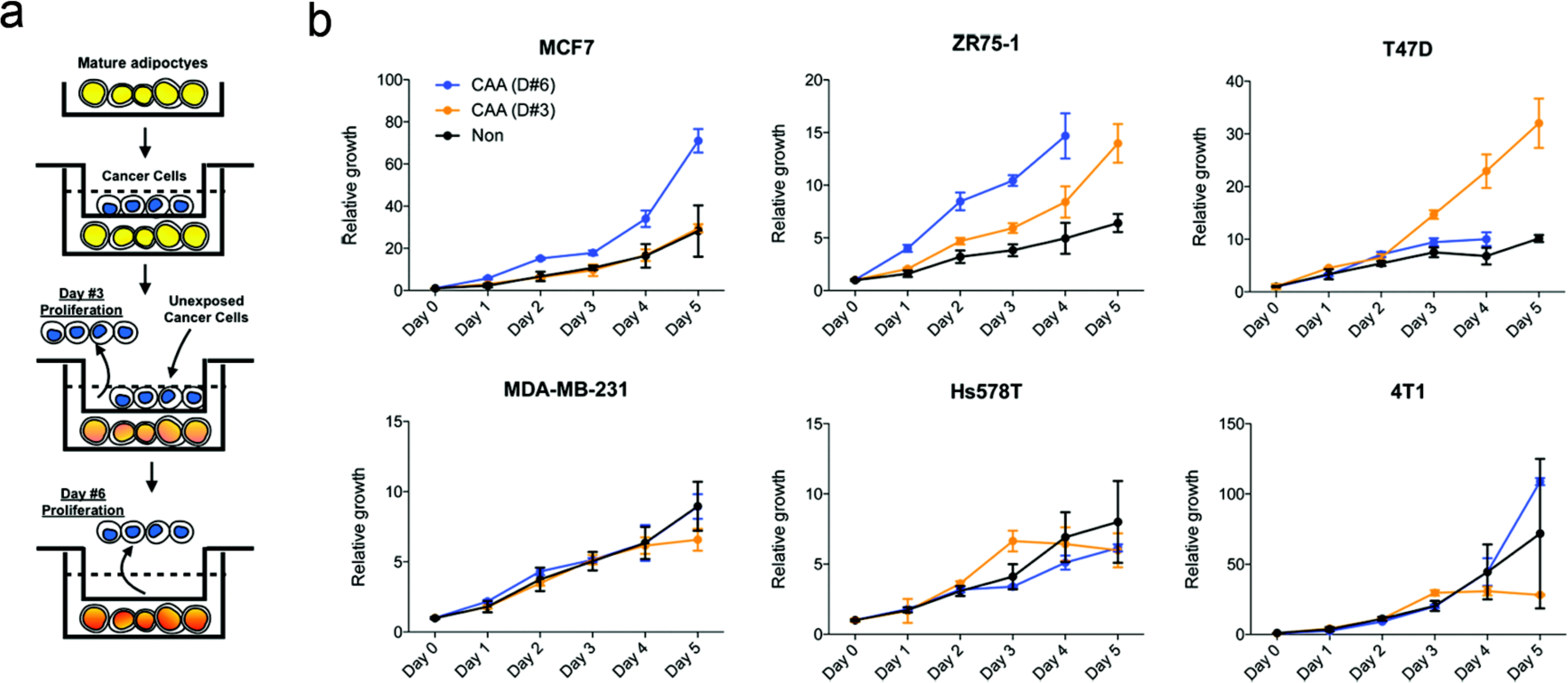
The effect of cancer-associated adipocytes (CAA) on proliferation of various breast cancer cells. The mature adipocytes were co-cultured with various breast cancer cells for three days, and the resulting CAAs were tested for the pro-proliferative effect on breast cancer cells (a). The degree of cell proliferation was measured by luminescent ATP quantification (b).

### Acquisition of inflammatory phenotype in CAAs

We analyzed transcriptional changes in adipocytes induced by the paracrine effect of breast cancer cell MCF7 and MDA-MB-231 by using indirect co-culture system. The genes involved in inflammation-related pathways, including inflammatory response, defense response, wound response, and chemotaxis, were significantly up-regulated in co-cultured adipocytes (Figure 4a). Additionally, genes involved in the peptidase inhibitor activity were also significantly changed. *Il6*, *Ptx3*, and *Timp1* were up-regulated genes with highest fold changes (Figure 4b), and the up-regulation of the genes was validated by PCR (Figure 4c). IL6 is suggested to be a major regulator of tumor-stroma interaction in cancer microenvironment [14], and Dirat et al [6] reported similar findings of increased IL6 expression in CAA.

**Figure 4.**
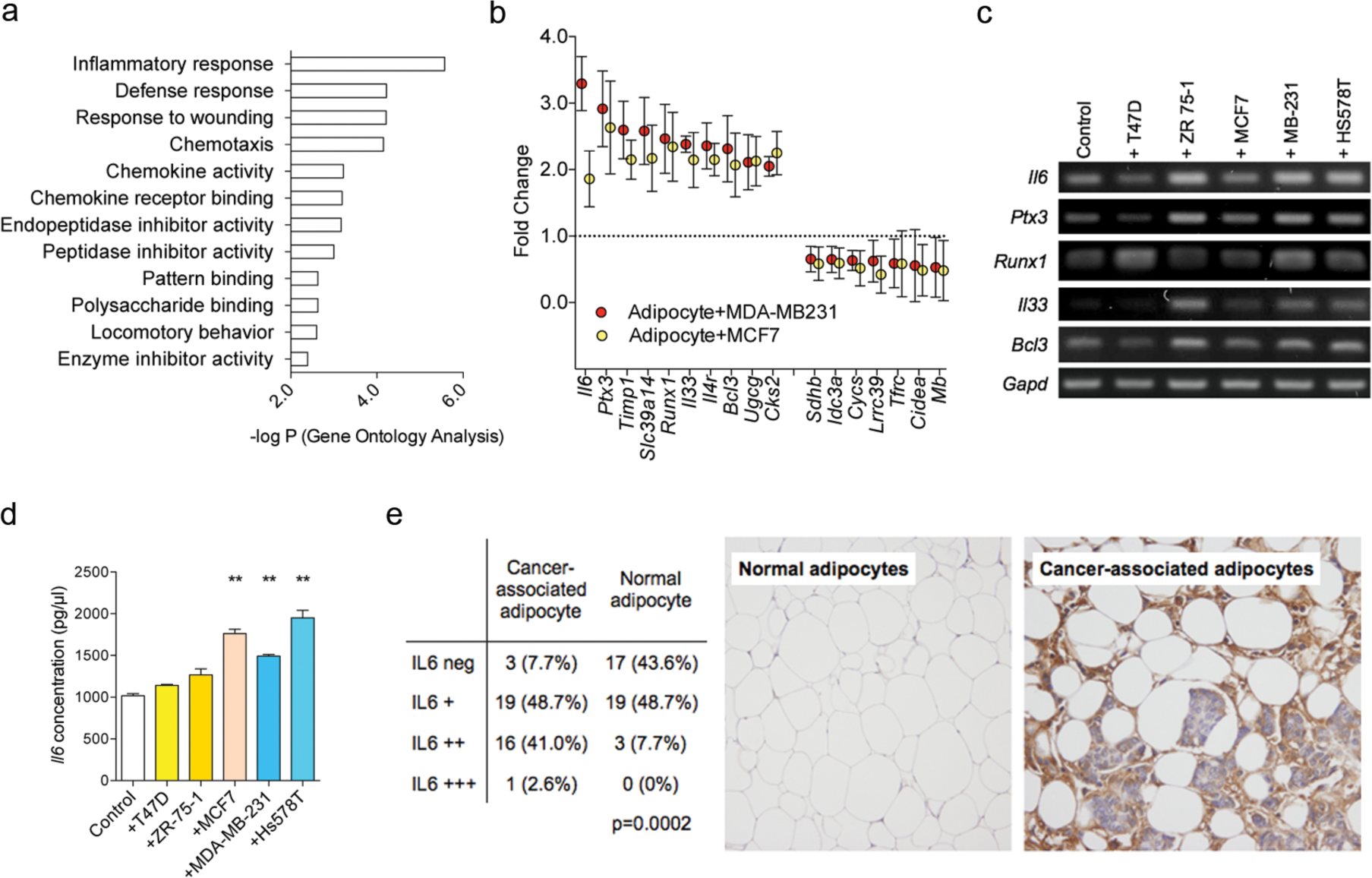
Gene expression profiles of cancer-associated adipocytes. Transcriptome analysis was done by using RNAs obtained from the indirect transwell co-culture system. Gene ontology analysis with DAVID software show that cancer-associated adipocytes are enriched with genes involved in inflammation and ECM remodeling (a). Differentially expression genes with highest fold change are shown in (b) and the differential expression was validated with PCR (c). ELISA against IL6 using secreted proteins from the cancer-associated adipocytes (e) and the IL6 expression levels in adipocytes from human breast cancer specimen (e) are shown.

We then assessed the IL6 protein levels in the media of co-cultured adipocytes and observed that the up regulation of Il6 gene resulted in increased secretion of IL6 (Figure 4d). To assess the IL6 expression levels in human breast cancer tissues, we examined the IL6 expression by immunohistochemistry using 39 consecutive breast cancer patients. Normal adipocytes, located far from the cancer cells, showed significantly lower frequencies of IL6 expression when compared to the adipocytes in the vicinity of breast cancer cells (Figure 4e). Our results suggest that breast cancer cells stimulate adjacent adipocytes via paracrine manner to express and secrete IL6 in breast cancer.

The expression levels of genes involved in the differentiation process of adipocytes were also explored using the gene expression data. While the co-cultured adipocyte did not show increased mesenchymal stem cell markers with the exception of *Cd44*, they showed significant down-regulation of the genes involved in the adipocyte differentiation (Supplementary Figure 2).

### mmu-miR-5112-mediated regulation of IL6 in CAA

MiRNAs can regulate the process of adipocyte differentiation, and are dysregulated in the various pathologic conditions involving adipose tissues.[15] The adipocytes co-cultured with MCF7 cells were profiled for their miRNA expression patterns to determine the differentially expressed miRNAs in CAA. Among the differentially expressed miRNAs, mmu-miR-5112 showed highest up-regulation in the co-cultured adipocytes (Figure 5a). The induction of miR-5112 expression in adipocytes by cancer cell was validated for MCF7 and MDA-MB-231 breast cancer cells (Figure 5b). The mmu-miR-5112 showed highest expression in the preadipocyte state and the expression was suppressed during the process of adipocyte differentiation (Figure 5c).

**Figure 5.**
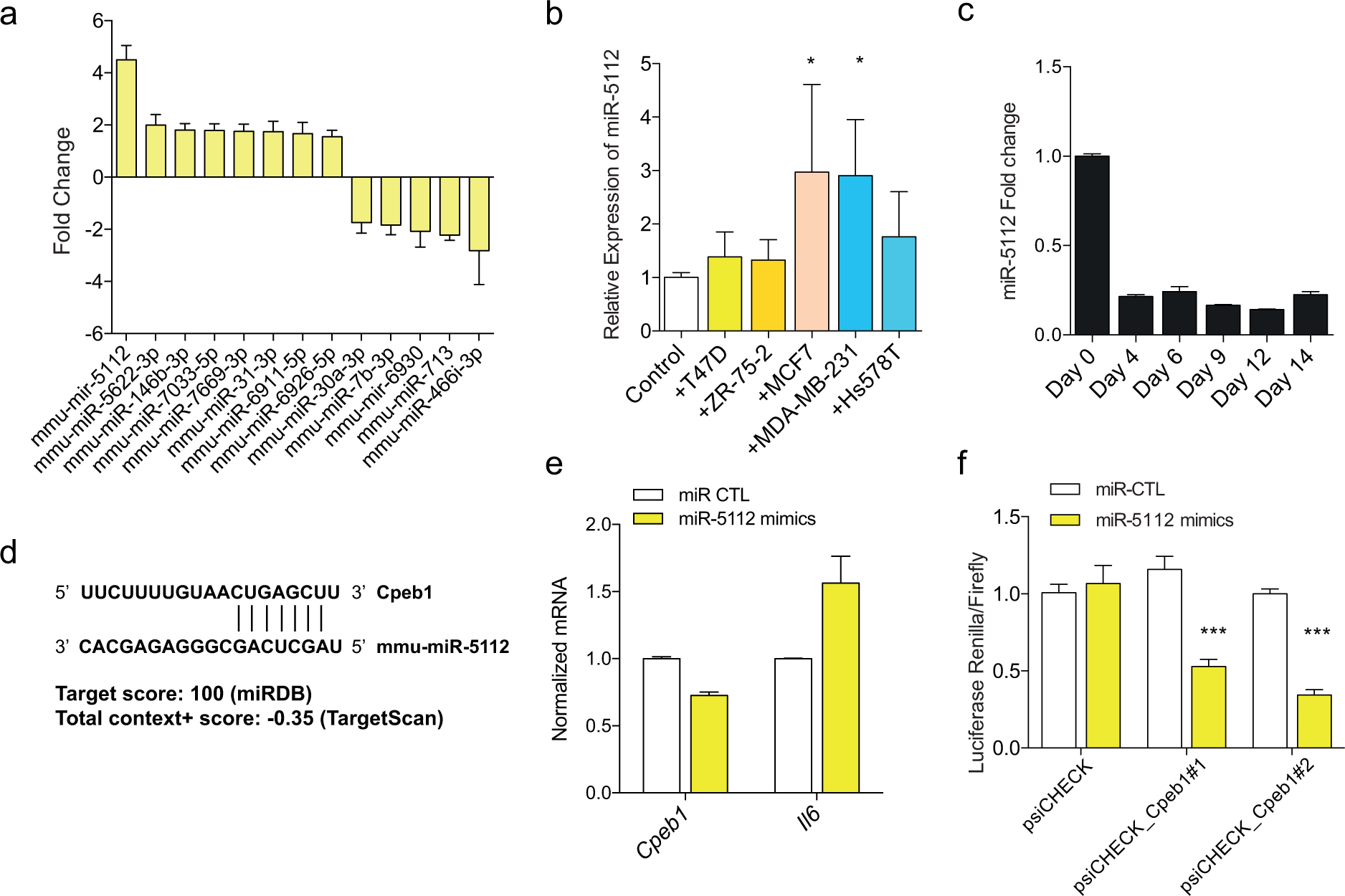
Regulatory role of mmu-miR-5112 in cancer-associated adipocyte (CAA) transition. microRNA profiles of CAA compared to normal adipocytes were determined by microarray (a). Expression of mmu-miR-5112 in adipocytes co-cultured with various cancer cells (b) and during the process of adipocyte differentiation (c). Predicted target site of Cpeb1 by mmu-miR-5112 (d) and the results of mmu-miR-5112 mimics treatment in Cpeb1 gene expression and translation measured by luciferase assay (e and f).

In silico analysis suggested Cpeb1 gene as a potential candidate for mmu-miR-5112 target gene (Figure 5d). CPEB1 gene is reported to be a negative regulator of IL6 production and an important regulator of inflammatory response.[16] To test this, we treated differentiated adipocytes with mmu-miR-5112 mimics and observed significant down-regulation of *Cpeb1* and increased level of *Il6* (Figure 5e). To confirm the regulatory role of mmu-miR-5112, the effect of mmu-miR-5112 on the translation activity in 3’ UTR of *Cpeb1* was measured by dual reporter luciferase assay. There was a significant decrease in the expression of luciferase reporter in cells transfected with mmu-miR-5112 mimics which demonstrates the suppression of Cpeb1 3’UTR translation by mmu-miR-5112 (Figure 5f). Taken together, our data suggest that the conversion of normal adipocytes into the inflammatory CAA is mediated by a miRNA-mRNA regulatory mechanism involving mmu-miR-5112-*Cpeb1*-*Il6*.

## DISCUSSION

In this study, we have analyzed the pathologic and molecular characteristics of CAA in human breast cancer specimens and determined the de-differentiated and inflammatory natures of CAA in breast cancer microenvironment. Additionally, by using indirect co-culture system for adipocytes and breast cancer cells, we propose a miRNA-based regulatory mechanism underlying the process of acquiring inflammatory phenotypes in CAA. Our results show that adipocytes acquire the morphologic and biologic characteristics of cellular de-differentiation process by adjacent breast cancer cells. Several studies have already reported the similar observation showing smaller cell size and down-regulation of adipogenic markers. [7,6,12,13,8] Studies have suggested that the de-differentiation process of CAA can be induced by up-regulation of ECM enzymes such as MMP11 [12] or cytokines such as TNFa or IL11 [13]. Indeed, our gene expression data shows that CAAs show significant upregulation of various cytokines and peptidase genes. The mRNA expression profiles of the CAA also showed dysregulation of genes involved in the pathways of inflammation, wound healing, and peptidase activities including *Il6* and *Ptx3*. IL6 carries diverse regulatory roles in breast cancer pathogenesis including remodeling the microenvironment, activation of EMT process, malignant transformation, and modulating breast cancer stem cell activities. [17-20]

The transformed adipocytes showed proliferation-promoting effect on breast cancer cells. However, the pro-proliferative effect of CAA was only seen in ER positive cell lines when tested in a panel of various types of breast cancer cells. Manabe et al [8] have also reported that adipocytes promote tumor growth in ER positive cells but not in ER negative MMT 060562 cells. These findings suggest that the tumor-promoting effect of adipocyte may depend on the molecular subtypes of breast cancer cells. The finding is in contrast to the work of Dirat et al [6] showing the pro-metastatic and invasive effect of CAA in both ER positive and negative cells. It is possible that the local growth promoting effect and the metastasis-promoting effect of adipocytes do not share similar mechanisms. Indeed, recent studies have suggested that growth promoting effect of microenvironmental cells on tumor mainly rely on local cellular interactions such as the exchange of metabolic supports or local production of aromatase enzymes. [5,21]

Based on the miRNA microarray and luciferase reporter assay, we propose a miRNA-based regulatory mechanism for the acquisition of inflammatory nature of CAA in breast cancer. Our data shows that mmu-miR-5112 is upregulated when the mature adipocytes are co-cultured with various breast cancer cells. This miRNA has been reported to suppress the *Cpeb1* gene that acts as a negative regulator of *Il6*.[16] Considering the pivotal roles of miRNAs in adipose tissue pathology as well as in the control of tumor microenvironment, it is likely that CAA transition in human adipocytes can be mimic our current observations in 3T3-L1 cells.[15,22] Similarly, Sung et al [23] have shown that let-7 can induce inflammatory stromal reaction in prostate cancer microenvironment.

There are several limitations in our study. First, we did not explore the functional effects of CAA in xenograft models. Additionally, we used mouse preadipocyte cells for our in vitro experiments, and therefore, the applicability of the present findings in human breast cancer microenvironment is not known. The in vivo effect of CAA in breast cancer progression and metastasis as well as the regulatory mechanisms in human adipocytes should be determined in further studies.

In conclusion, our results describe a potential microRNA-based regulatory mechanism underlying the transition between normal adipocytes and inflammatory/de-differentiated CAA in breast cancer. Furthermore, we show that the pro-survival effect of the CAA can be breast cancer subtype-specific. Additional studies to address the molecular links that explain the subtype-specific effect may provide a window for microenvironment-targeted therapies.

## CONFLICTS OF INTERESTS

Authors declare no conflict of interests

## ACKNOWLEDGMENT AND FUNDING INFORMATION

This work was supported by the Research Resettlement Fund for the New Faculty of Seoul National University, Basic Science Research Program through the National Research Foundation of Korea (NRF) funded by the Ministry of Education, Science and Technology (2012R1A1A2005929 and 2015R1D1A1A02061904), by the grant from the National R&D Program for Cancer Control, Ministry for Health and Welfare, Republic of Korea (A1520250), and by the grant of the Korea Health Industry Development Institute (KHIDI), funded by the Ministry of Health & Welfare, Republic of Korea (HI14C1277 and HI13C2148).

## FIGURE LEGENDS

**Supplementary Figure 1.**
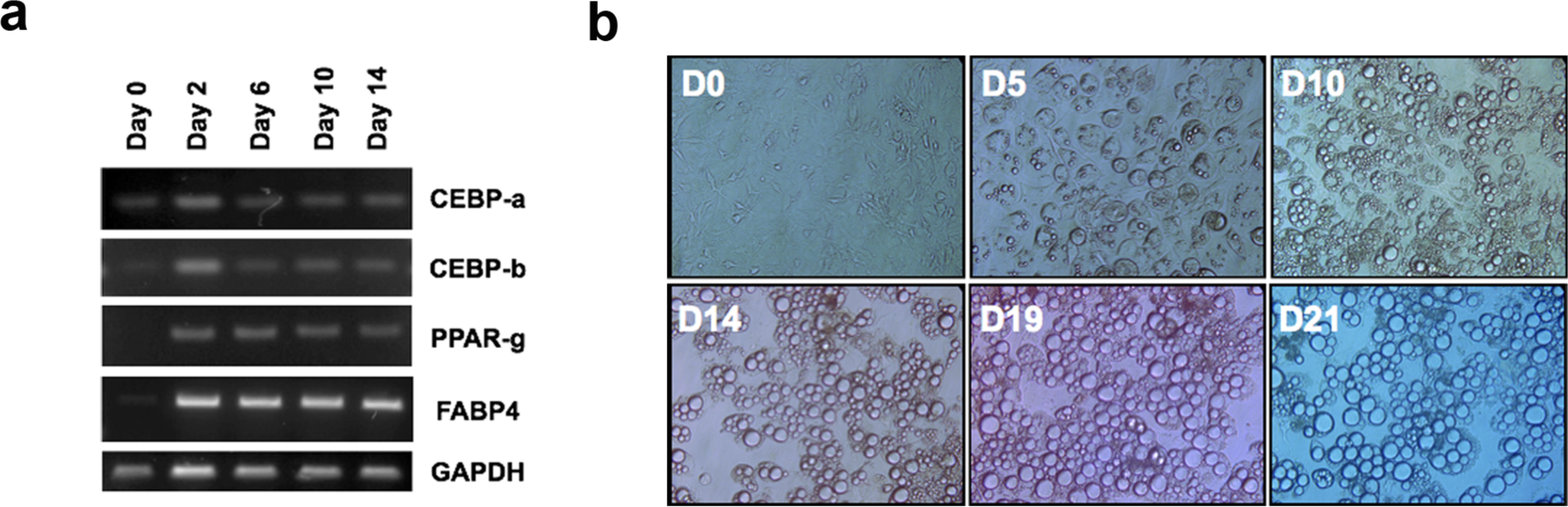
The changes in the adipocyte differentiation-related genes and microscopic shapes in NIH-3T3-L1 cells during adipogenic differentiation. Along with the adipogenic gene expression (a), the cells showed increased accumulation of lipid drops and increase in the cell sizes (b).

**Supplementary figure 2.**
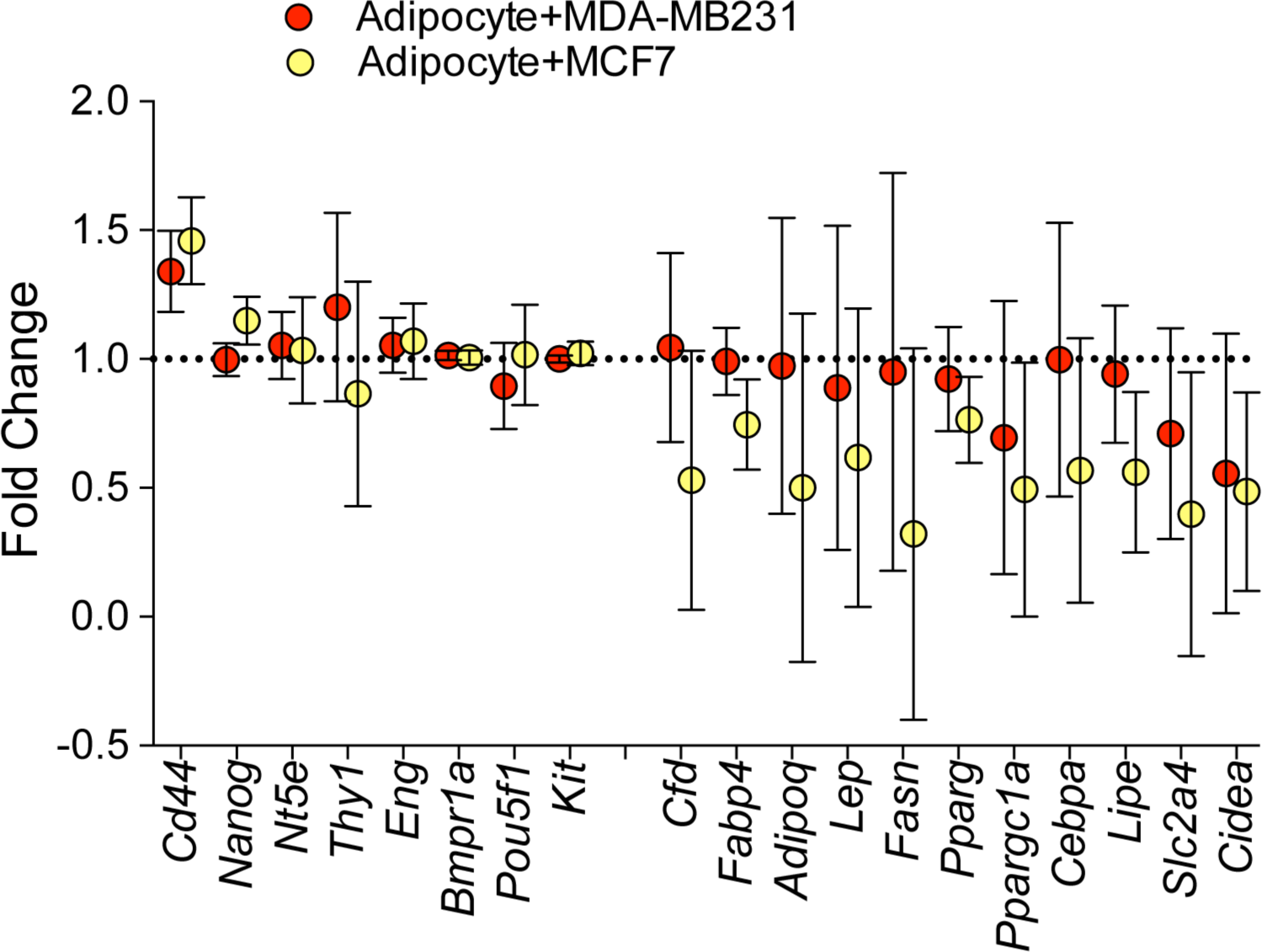
Expression levels of differentiation markers in the gene expression data. Genes in red and blue represent the genes expressed in mesenchymal stem cells and in differentiated adipocytes, respectively.

